# Aging shapes infection profiles of influenza A virus and SARS-CoV-2 in human lung slices

**DOI:** 10.1101/2024.04.14.589423

**Authors:** Melanie Brügger, Carlos Machahua, Beatrice Zumkehr, Christiana Cismaru, Damian Jandrasits, Patrick Dorn, Thomas M. Marti, Gert Zimmer, Volker Thiel, Manuela Funke-Chambour, Marco P. Alves

## Abstract

The recent coronavirus disease 2019 (COVID-19) outbreak revealed the susceptibility of elderly patients to respiratory virus infections, showing cell senescence or subclinical persistent inflammatory profiles and favouring the development of severe pneumonia. In our study, we evaluated the potential influence of lung aging on the efficiency of replication of influenza A virus (IAV) and severe acute respiratory syndrome coronavirus 2 (SARS-CoV-2), as well as determined the pro-inflammatory and antiviral responses of the distal lung tissue. Using precision-cut lung slices (PCLS) from donors of different ages, we found that pandemic H1N1 and avian H5N1 IAV replicated in the lung parenchyma with high efficacy. In contrast to these IAV strains, SARS-CoV-2 early isolate and Delta variant of concern (VOC) replicated less efficiently in PCLS. Interestingly, both viruses showed reduced replication in PCLS from older compared to younger donors, suggesting that aged lung tissue represents a sub-optimal environment for viral replication. Regardless of the age-dependent viral loads, PCLS responded to infection with both viruses by an induction of IL-6 and IP-10/CXCL10 mRNAs, being highest for H5N1. Finally, while SARS-CoV-2 infection was not causing detectable cell death, IAV infection caused significant cytotoxicity and induced significant early interferon responses. In summary, our findings suggest that aged lung tissue might not favour viral dissemination, pointing to a determinant role of dysregulated immune mechanisms in the development of severe disease.

**New & Noteworthy:** PCLS from donors of varying ages were exposed to SARS-CoV-2 or IAV. Notably, the latter exhibited the highest replication efficacy, triggering early interferon responses, elevated IL-6 and IP-10/CXCL10 mRNAs expression, and significant cell death compared to SARS-CoV-2. Overall, across all age groups, the pulmonary environment showed sustained immunocompetence. For both viruses, older donor-derived PCLS displayed reduced viral permissiveness, suggesting aged lung tissue might not favour viral dissemination, implying other factors contribute to severe disease development.

## INTRODUCTION

Viral pneumonia induced by influenza A virus (IAV) and severe acute respiratory syndrome coronavirus 2 (SARS-CoV-2) is a leading cause of death from infectious diseases worldwide (1–3). Importantly, there is a profound difference for the clinical severity of coronavirus disease 2019 (COVID-19) depending on age. Specifically, while advanced age is associated with higher risk for severe disease and death, children are rarely affected, and severe pneumonia is unusual in this population (4). On the other hand, infection by IAV is associated with increased mortality in older patients, but also pose a risk for severe disease in young children (5–7).

The increased susceptibility of the elderly to severe respiratory virus infections is due to a combination of factors associated with aging, including comorbidities, a decline in lung function, cellular senescence, and dysregulated immunity. This includes impairment of local innate immune mechanisms, an hyperinflammatory phenotype related to age called “inflammaging”, and a decline in humoral and cellular responses, termed “immunosenescence” (8–10). Infection with a number of viruses, such as measles virus, human respiratory syncytial virus, and coronaviruses, can induce premature cellular senescence (11). Interestingly, infection of lung tissue by IAV induced premature aging as indicated by the appearance of senescent cells and subsequently, infection of senescent cells resulted in increased viral replication (12). Similarly, SARS-CoV-2 was shown to induce senescence through multiple mechanisms with the potential to trigger COVID-19-related “cytokine storm” and organ damage (13).

Nevertheless, the impact of age and the local pulmonary microenvironment on acute respiratory virus infection has not been elucidated so far, mainly due to experimental limitations and because no suitable animal model is available. In previous studies, lung explants (14–18) and lung slices (19) have been applied to study IAV infection and virulence, showing the high efficiency of virus replication in cultured lung tissue from the lower respiratory tract. Lung explants have also been used to study the replication potential of SARS-CoV-2 variants, the cellular tropism of SARS-CoV and SARS-CoV-2, and to elaborate subsequent immune responses (17, 18, 20). However, these studies did not investigate the impact of the tissue donors’ age. Of note, we and others have demonstrated an age-dependent susceptibility of the upper respiratory epithelium towards respiratory virus infection (21, 22). Thus, we hypothesized that the features of the lung from older individuals, particularly the alveolar epithelium of the distal lung, may influence viral replication and contribute to the adverse outcomes during IAV or SARS-CoV-2 infections in the elderly. We employed a state-of-the-art *ex vivo* model of respiratory virus infection based on organotypic cultures of lung parenchymal tissue, namely precision-cut lung slices (PCLS) (23–26). This culture model preserves the bronchiolar and alveolar structures with minimal changes during culture for up to two weeks (27), allowing the study of cell behaviour and cell-cell interactions in their native environment (26). Due to these characteristics, PCLS are bridging the gap between simple *in vitro* and complex *in vivo* systems (24), by recapitulating pathophysiological conditions and local inflammatory responses, that are believed to occur *in vivo* (28). The advantage of PCLS compared to lung explants is to ensure homogeneity and reproducibility, thus reducing the degree of variability between donors (29, 30).

In this study, we investigated IAV and SARS-CoV-2 replication and pro-inflammatory and antiviral responses induced by infection of “older” *versus* “younger” distal lung tissue. We used human PCLS, obtained from several donors with different ages, for viral infection studies using an early isolate of SARS-CoV-2 and an isolate of the Delta VOC, as well as the 2009 pandemic IAV H1N1 and avian IAV H5N1.

## MATERIALS AND METHODS

### Biosafety statement

All experiments with infectious H5N1/A/turkey/Turkey/1/2005 and SARS-CoV-2 were performed in an enhanced biosafety level 3 (BSL3) containment laboratory at the Institute of Virology and Immunology, Mittelhäusern, Switzerland, which followed the approved standard operating procedures of BSL3 facility of relevant authorities in Switzerland (authorization no. A202819). All the personnel involved in these experiments received advanced training and wore appropriate personal protective equipment, including Tyvek overall and hood, and a Jupiter or Versaflo respirator with an HEPA filter system (all 3M).

### Cell lines and culture conditions

Madin-Darby canine kidney type II (MDCK-II) cells were kindly given by Georg Herrler (University of Veterinary Medicine Hannover, Germany) and were cultured in MEM (Thermo Fisher Scientific) supplemented with 5% fetal bovine serum (FBS). Human embryonic kidney 293T (HEK-293T) cells (American Type Culture Collection (ATCC), Manassas, USA; CRL-3216), were maintained in DMEM supplemented with 10% FBS. VeroE6 cells were kindly provided by Doreen Muth, Marcel Müller, and Christian Drosten (Charité, Berlin, Germany) and cultured in DMEM GlutaMax supplemented with 10% FBS and 1% HEPES (all Thermo Fisher Scientific). VeroE6-TMPRSS2 cells (NIBSC Research Reagent Depository, UK) were cultured in DMEM GlutaMAX with 10% FBS and 1X non-essential amino acids (Thermo Fisher Scientific). When passaging, 200 µg/mL of geneticin (G418) were added to the culture medium. All cells were maintained in a humidified incubator at 37°C, 5% CO_2_.

### Human precision-cut lung slice cultures

Human PCLS were prepared as previously reported by us (31, 32). Briefly, lung tissue from the edges of tumour resections, with no signs of inflammation or emphysema, were collected at the Inselspital, Bern University Hospital (approved by Swiss Ethics Commission, license: KEK-BE_2018-01801). All lung tissue resections were tested for SARS-CoV-2 by RT-qPCR. Once in the lab, tissues were processed within six hours post-surgery: 1 cm^3^ low-melting agarose-perfused (2%) lung tissues were cut in 400 μm slices in the Compresstome VF-310-0Z Vibrating Microtome (Precisionary), following the manufacturer’s recommendations. Slices were placed in 12-well plates with DMEM GlutaMax medium, 1% FBS, HEPES 20mM, and 1X Antibiotic-Antimycotic (Thermo Fisher). PCLS were incubated at 37°C, 5% CO_2_ in an humidified incubator, and medium was changed every day for one to seven days prior infection.

### Human well-differentiated primary nasal epithelial cells

Generation of human well-differentiated nasal epithelial cell cultures (WD-NECs) was done as previously described (31). In short, primary NECs were obtained commercially (MucilAir, Epithelix Sàrl, Geneva, Switzerland). Cells were seeded and maintained on 24-well transwell inserts (Corning) at the air-liquid interface in MucilAir culture medium (Epithelix Sàrl, Geneva, Switzerland) in a humidified incubator at 37°C, 5% CO_2_. Culture medium was changed every three days.

### Viruses

The eight-plasmid pHW2000 reverse genetics system of H1N1 A/Hamburg/4/2009 (GenBank accession nos.: GQ166207, GQ166209, GQ166211, GQ166213, GQ166215, GQ166217, GQ166219, GQ166221) was provided by Hans-Dieter Klenk (University of Marburg, Marburg, Germany). For generation of infectious virus, 10^6^ MDCK-II cells and 2x10^6^ HEK-293T cells were seeded into 10 cm cell culture dishes and grown overnight at 37°C, 5% CO_2_. The co-culture was transfected with the 8-plasmid set (2 μg of each plasmid) using Lipofectamine 2000 transfection reagent (Fisher Scientific). At 24 h post transfection, the cells were washed with PBS and maintained in serum-free medium supplemented with 0.2% (w/v) of bovine serum albumin (BSA), 1% (v/v) of penicillin/streptomycin, and 1 μg/ml of acetylated trypsin. Passage 0 was obtained six days post-transfection, when the cell medium was collected, FBS was added to a final concentration of 5% and cell debris was removed by centrifugation (3000 rpm, 10 min, 4°C). Two passages on MDCK-II, in the presence of trypsin, were performed to obtain the working stock, which was titrated on MDCK-II cells, as described below, before use. H5N1 A/turkey/Turkey/2005 was kindly provided by Dr. W. Dundon, (IZSV Instituto Zooprofilattico Sperimentale delle Venezie, Venice, Italy). Virus was passaged twice in eggs and allantoic fluid was clarified using low-speed centrifugation as described previously (33). A SARS-CoV-2 early isolate (SARS-CoV-2/München1.1/2020/929, “Early isolate” in subsequent text and figures), kindly provided by Daniela Niemeyer, Marcel Müller, and Christian Drosten (Charité, Berlin, Germany), was passaged once in VeroE6 cells. To produce the working stock, SARS-CoV-2 Early isolate was then propagated once more in VeroE6 cells at a multiplicity of infection (MOI) of 0.01 plaque forming unit (PFU) per cell. Therefore, the respective volume of the original isolate was added to the cells in DMEM GlutaMax supplemented with 10% FBS and incubated for 36 h at 37°C, 5% CO_2_. SARS-CoV-2 Delta isolate (SARS-CoV-2 Delta AY.127 (hCoV-19/Switzerland/BE-IFIK-918-4879/2021, L5109, passage of EPI_ISL_1760647) (34) was passaged three to four times accordingly on VeroE6-TMPRSS2 cells. Next, the supernatant was centrifuged to eliminate cell debris, and then stored in aliquots at −70°C.

### Infection of human precision-cut lung slice cultures

PCLS were infected one to seven days post-surgery and experiments were conducted within four to ten days post-surgery. Similarly-sized PCLS were infected with either Mock or virus at 10^6^ PFU of IAV H5N1/A/turkey/Turkey/2005 and H1N1/Hamburg/4/2009 or SARS-CoV-2 Early and Delta isolate, in 750 µL infection medium (DMEM GlutaMAX, supplemented with 0.1% FBS, 100 units/mL of penicillin and 100 μg/mL streptomycin, and 2.5 μg/mL of Amphotericin), for 2 h at 37°C, 5% CO_2_. Afterwards, PCLS were washed twice with PBS with Ca^2+^ and Mg^2+^, to remove residual virus inoculum and then fresh growth medium (DMEM GlutaMAX, supplemented with 1% FBS, 100 units/mL of penicillin and 100 μg/mL streptomycin, and 2.5 μg/mL of Amphotericin) was added. Every 24 h medium was changed and 4, 24, 48, and 72 h post-infection (h p.i.), PCLS were collected for RNA extraction in ice-cold TRIzol reagent (Thermo Fisher) or washed twice in PBS, transferred to embedding cassettes, and then fixed in 4% formalin for 24 h until further processing for immunohistochemistry and immunofluorescence staining. Supernatants were frozen at -70°C until further processing.

### RNA isolation and quantitative PCR

RNA isolation of PCLS was performed by the combination of two procedures, as previously described (32). PCLS were placed into MagNA Lyser Green Beads tube (Roche Diagnostics) with 700 μl ice-cold TRIzol reagent and disrupted in a Bullet Blender Tissue Homogenizer (Next Advance). After the separation of phenol-chloroform, approx. 250 µl aqueous phase was mixed 1:1 with 75% ethanol and transferred into the Zymo-Spin IC Column from the RNA Clean & Concentrator Kit (Zymo Research). RNA clean-up and collection was done following the manufacturer’s instructions.

RNA of WD-NECs was extracted using the Nucleospin RNA Plus Kit (Macherey-Nagel) according to manufacturer’s protocol. For the detection of viral RNA (except H1N1) and host mRNA, the Omniscript RT kit (Qiagen) using random hexamers (Thermo Fisher Scientific) was used for reverse transcription and synthesis of complementary DNA (cDNA). Quantitative PCR (qPCR) was performed on an ABI Fast 7500 Sequence Detection System (Applied Biosystems) with target-specific primers, using the TaqMan Gene Expression Assay (Applied Biosystems) or the Fast SYBR Green Assay (Thermo Fisher Scientific). For H1N1/A/Hamburg/4/2009, one-step cDNA synthesis and qPCR were performed on a 7500 Real-Time PCR System (Thermo Fisher Scientific) using the AgPath-ID One-Step RT-PCR kit (Thermo Fisher Scientific) with target-specific primers. Results were analysed using the SDS software (Applied Biosystems), whereas relative expression was calculated with the ΔΔCT method as described (35). Expression levels were normalized to the housekeeping 18S ribosomal RNA (rRNA) or beta-2-microglobuline (B2M) for the analysis of the genes of interest (32, 36–45). The sequences and concentrations of primers and probes used are listed in **Table 1**. For H5N1 A/turkey/Turkey/2005, the Flu panA mod primer and probes were used and for H1N1/Hamburg/4/2009, they were combined with the Flu M swine reverse primer (41).

**Table 1.**
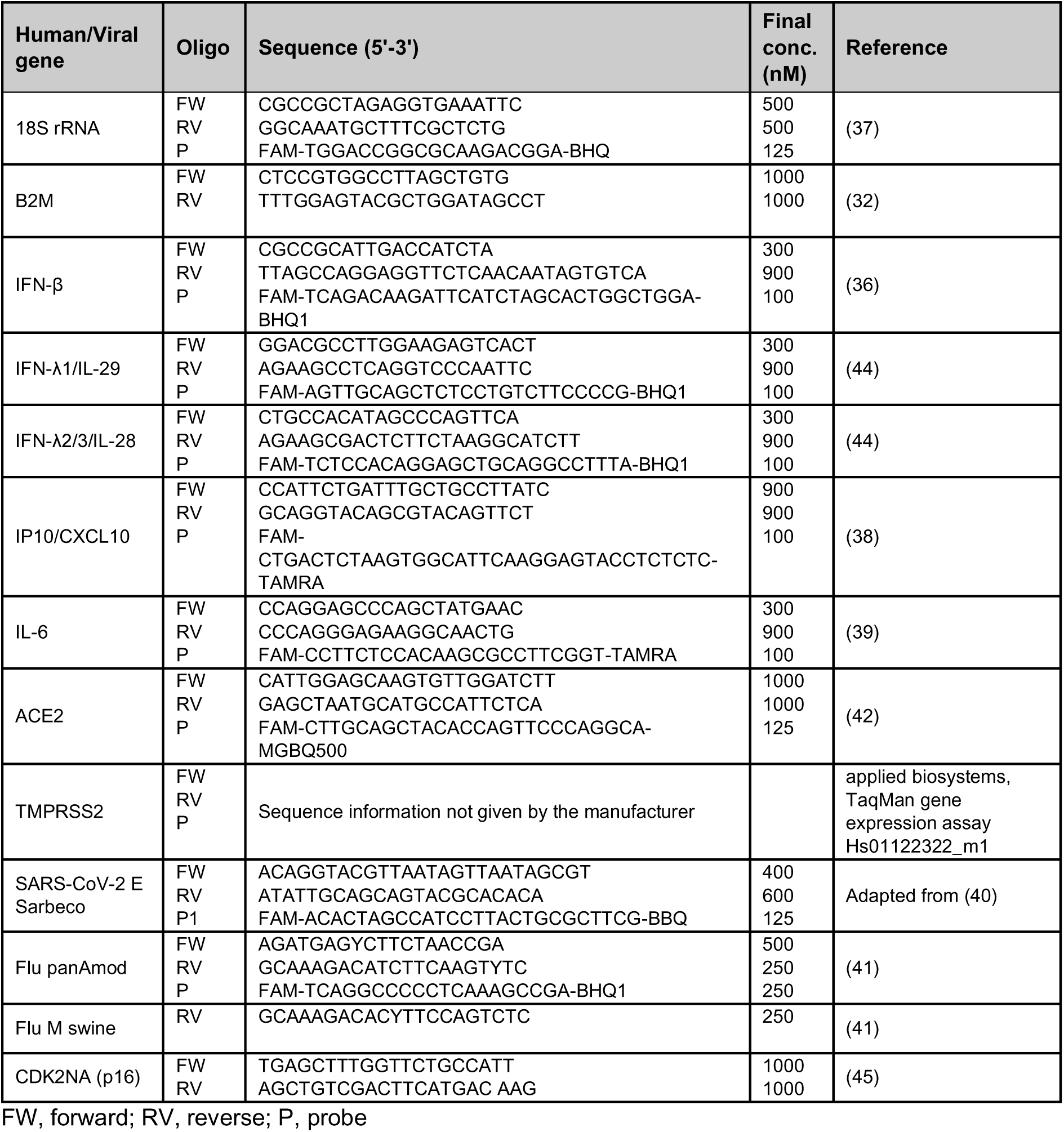
List of primer and probe sequences and concentrations used.

### Infectious virus titration

For IAV titration, MDCK-II cells were seeded at 10^5^ cells/well density into 96-well plates and cultured overnight before infection. The virus samples were 10-fold serial-diluted in serum-free growth medium, and each sample was titrated in duplicate (40 µl inoculum/well). After a 90-minute adsorption at 37°C, an overlay containing MEM with 1% (w/v) methylcellulose, 2% FBS and 1% (v/v) penicillin/streptomycin solution were added to each well (160 µl). 24 h p.i. cells were washed twice with pre-warmed PBS and fixed in 4% formalin for 15 min. This was followed by 0.1 M glycine-quench washes, and a 5-min permeabilization with 0.25% (v/v) of Triton X-100 in PBS. Primary antibody staining against viral nucleoprotein (NP) was performed using the mouse monoclonal HB-65 (ATCC, clone H16-L10-4R5) antibody diluted 1:50 in PBS, followed by the secondary antibody goat anti-mouse IgG conjugated to Alexa Fluor 488 (ThermoFisher), diluted 1:500 in PBS. Both antibodies were incubated for 1 h at 21°C and were each followed by three washing steps with PBS. The infected cell foci were counted manually using the Observer.Z1 fluorescence microscope (Zeiss, Feldbach, Switzerland) using a 10x magnification objective, and based on this the virus titers were calculated as PFU/ml. For SARS-CoV-2 titration, VeroE6 cells were seeded at 3x10^6^ cells per 24-well plate and cultured overnight before infection. The samples were diluted 1:2 and in a 1:10-fold serial dilution and added to the wells. After 1 h of incubation at 37°C, 5% CO_2_, the medium was removed and cells were overlayed with a 1:1 mix of DMEM, 20% FBS, 2X HEPES, 2X Pen/Strep and 2.4% Avicel. 48-72 h later, the overlay was removed, cells were washed with PBS, fixed in 4% formalin for 15 min, and stained with crystal violet for another 15 min. Then, wells were washed with tap water and dried before plaques were counted. Virus titers were calculated as PFU/ml.

### Immunohistochemistry staining of SARS-CoV-2 infected PCLS

Formalin-fixed PCLS were embedded in paraffin using a standard protocol by the COMparative PATHology platform (COMPATH) at the Institute of Pathology and Institute of Animal Pathology, University of Bern, Switzerland. Briefly, PCLS were dehydrated in an ethanol series (70%, 96%, 100%, Grogg), cleared with xylol (Sigma-Aldrich), and embedded in paraffin. Using a Leica RM2135 microtome (Leica Biosystems), 7 µm thick sections were cut and mounted on glass slides (Fisherbrand Superfrost Plus, Fisher Scientific). For dewaxing, the slides were incubated at 60°C for 1 h, followed by rehydration with three times xylol and a series of descending concentrations of ethanol (2x 100%, 2x 95%, 80% to 70%). For immunohistochemistry, endogenous peroxidase activity was inhibited by 3.25% H_2_O_2_ in 70% methanol for 10 min at room temperature followed by washing the slides in PBS. For antigen retrieval, the slides were then boiled using a microwave in 10 mM citrate buffer (pH 6.0) for two times 5 min. After washing in PBS, samples were incubated in PBS with 1% BSA (Sigma-Aldrich) for 30 min at room temperature to block nonspecific antibody binding. Then, primary antibody targeting the nucleocapsid (N) protein of SARS-CoV-2 (Rockland, 1:2’500, rabbit polyclonal IgG) in PBS with 1% BSA was incubated over night at 4°C in a humidified chamber. After washing, slides were incubated with biotinylated goat anti-rabbit (DAKO), followed by HRP-conjugated streptavidin (DAKO), both for 10 min at room temperature protected from light, with an interposed washing step. Next, slides were washed in PBS and AEC-substrate chromogen was incubated for 10 to 15 min at room temperature. Then, slides were washed in tap water, followed by distilled (d)H_2_O and haematoxylin staining of the nuclei for 2 min. Afterwards, slides were again washed in tap water followed with dH_2_O. Slides were then covered with Aquatex mounting media (Merck) and a coverslip.

### Immunofluorescence staining of PCLS

After 24 h, formalin-fixed PCLS were washed twice with PBS and stored in 70% ethanol at 4°C, until further processing. For staining, the ethanol was removed by three 10-min washes in PBS at room temperature. Permeabilization was done with PBS-Triton X-100 0.3% supplemented with 2% BSA for 15 min at room temperature. Blocking was then performed with PBS-Triton X-100 0.3% supplemented with 30 mg/ml milk powder and 10% FBS (blocking solution), for 1 h at room temperature. Primary antibody against advanced glycosylation end-product specific receptor (AGER, R&D Systems, 1:250), pro-surfactant protein C (pro-SFTPC, Novus Biologicals, 1:800), IAV nucleoprotein (NP, 1:50), and SARS-CoV-2 nucleocapsid protein (N, clone E16C, Fisher Scientific, 1:25) were diluted in blocking solution and then added and incubated overnight at 4°C. This was followed by three 10-min washes in PBS-Triton 0.1% at room temperature. Finally, the secondary antibody donkey anti-goat IgG (1:400, AGER) or goat anti-rabbit IgG (1:400, pro-SFTPC), both conjugated to Alexa Fluor 546, goat anti-mouse IgG (1:500, IAV) or goat anti-mouse IgG2b (1:250, SARS-CoV-2), both conjugated to Alexa Fluor 488, Phalloidin conjugated to Alexa Fluor 546 (1:100, for IAV staining only), and DAPI staining at 1:1000 dilution, in blocking solution was added to the PCLS and incubated for 5 h at 4°C. The samples were mounted on glass slides using the EMS Shield Mount with DABCO™ (AGER and pro-SFTPC, EMS, USA), ProLong Gold antifade reagent (IAV, Fisher Scientific), or Mowiol (SARS-CoV-2, Merck/Calbiochem). All the incubations and washing steps were performed under soft rocking for better results. The imaging was performed using the Nikon confocal A1 microscope (IAV NP, Phalloidin) or by laser scanning confocal microscopy using Olympus FV3000 with FluoView FV3000 system software (SARS-CoV-2 N, AGER, pro-SFTPC) and Zeiss LSM 710 (AGER, pro-SFTPC). Images were processed and rendered in the Nikon NIS Elements Confocal version 4.30 software (IAV) and Fiji, or ImageJ image processing package v1.54f or v1.53q, respectively (SARS-CoV-2, AGER, pro-SFTPC).

### Cytotoxicity assay

Lactate dehydrogenase (LDH) release was assessed in the supernatants (four technical replicates per sample) of Mock-, SARS-CoV-2-, and IAV-infected PCLS using the CytoTox 96 Non-Radioactive Cytotoxicity Assay (Promega) following manufacturer’s instructions. Absorbance was measured at 490 nm with the Biotek 800 TS absorbance reader (Agilent technologies). To calculate the percentage of cytotoxicity, growth medium background was subtracted and resulting values were divided by the maximum LDH release, which was obtained by taking the supernatant of adherent HEp-2 cells (ATCC, CCL-23), incubated with 1X lysis solution (provided by the manufacturer) for 30 min.

### Statistical analysis

Statistical analysis was performed using GraphPad Prism v9 and v10. The specific statistical tests used are indicated in the figure legends.

## RESULTS

### PCLS cultures from aged lungs present a senescent phenotype

To investigate the impact of aging on respiratory virus infection, lung tissue from donors with ages ranging from 36 to 81 years were used to generate PCLS cultures (**Fig. 1A and Table 2).** PCLS retained the native microstructure and cellular composition of the alveolar epithelium, as confirmed by staining for the advanced glycosylation end-product specific receptor (AGER), which is expressed at high baseline levels primarily in alveolar type 1 cells (46), and for pro-surfactant protein C (SFTPC), a marker for alveolar type 2 cells **(Fig. 1B, C).** Further, we observed a significant association between tissue donor age and mRNA levels of the senescence marker *CDKN2A* (p16), indicating higher numbers of senescent cells in PCLS obtained from older donors **(Fig. 1D)**.

**Figure 1.**
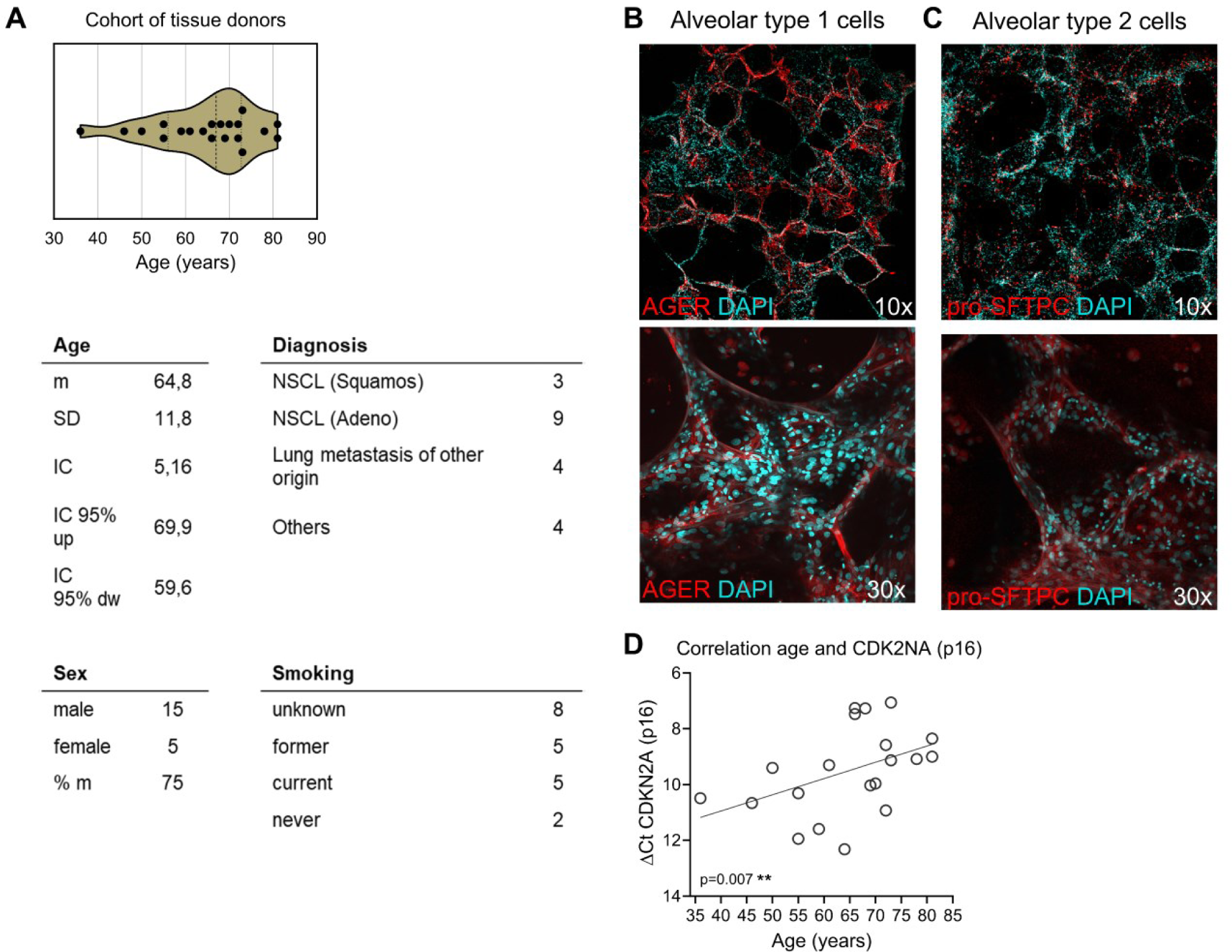
PCLS cultures from aged lungs present a senescent phenotype. **(A)** Violin plot representing the age distribution of the individual donors used for the generation of the PCLS cultures. The dashed vertical line indicates the median value, whereas the dotted lines indicate the interquartile ranges. Each symbol represents an individual donor. The table summarizes patient’s characteristics: age (m, SD), sex (% male, m), diagnosis and smoking status (n). **(B, C)** Representative confocal microscopy evaluation of PCLS stained for **(B)** AGER, highly expressed in alveolar type 1 cells, or **(C)** the alveolar type 2 cell marker pro-SFTPC. Micrographs are projections of 3D captures. Magnification 10X and 30X. AGER and pro-SFTPC, red; DAPI, light blue. **(D)** Correlation of deltaCT CDKN2A (p16) and donor age. Association was tested using the one-tailed Spearman rank correlation test. Each symbol represents an individual donor.

**Table 2.** Characteristics of the pulmonary tissue donors.

| Patient | Birth | Age | Sex | Smoking status | Diagnosis |
| --- | --- | --- | --- | --- | --- |
| BE1043N | 1940 | 81 | male | unknown | NSCL (Squamous) |
| BE1050N | 1949 | 72 | male | unknown | NSCLC (Adeno) |
| BE1104N | 1955 | 66 | male | 15 py, former smoker | NSCLC (Adeno) |
| BE1128N | 1971 | 50 | female | most probably never smoker | Lung metastasis of Melanoma |
| BE1133N | 1966 | 55 | male | 50 py, smoker | NSCLC (Adeno) |
| BE1144N | 1953 | 68 | female | unknown | NSCLC (Adeno) |
| BE1150N | 1951 | 70 | male | unknown | Infectious (Pneumonia) |
| BE1163N | 1967 | 55 | male | 35py, smoker | benign (lymphcytic nodule) |
| BE1164N | 1963 | 59 | male | ? py, current smoker | Infectious (Pneumonia) |
| BE1165N | 1976 | 46 | male | 25py, current smoker | Lung metastasis of carcinoma of testicle |
| BE1166N | 1953 | 69 | male | 30py, former smoker | NSCLC (Squamous) |
| BE1203N | 1956 | 66 | male | 30 py | NSCLC (Adeno, SMARC-A4-deficiency) |
| BE1205N | 1961 | 61 | male | 22py, stopped 2018 | NSCLC (Adeno) |
| BE1207N | 1949 | 73 | male | 60py, stopped 2021, now e-cigarettes | NSCLC (Squamous) |
| BE1261N | 1987 | 36 | male | unknown | Lung metastasis of colon carcinoma |
| BE1282N | 1942 | 81 | male | never smoker | NSCLC (Adeno) |
| BE1288N | 1951 | 72 | female | unknown | Atypical carcinoid |
| BE1290N | 1959 | 64 | female | unknown | NSCLC (Adeno) |
| BE1292N | 1950 | 73 | female | 20 py, stopped 1996 | NSCLC (Adeno) |
| BE1294N | 1945 | 78 | male | unknown | Lung metastasis of Melanoma |

NSCLS, Non-small cell lung cancer; py, pack-year index

### Human PCLS are productively infected by pandemic and avian IAV strains

We first studied the effect of age on IAV infection using pandemic H1N1 and avain H5N1 strains. Specifically, we infected PCLS from three and four donors (64, 72, 81 and 36, 73, 78, 81 years old, respectively) with 10^6^ PFU of A/Hamburg/4/2009 (H1N1) or A/turkey/Turkey/2005 (H5N1), respectively. Two h p.i., PCLS were washed twice and incubated at 37°C for 4 to 72 h. To preserve tissue viability, culture medium was changed every day, and at selected time points p.i., individual PCLS from the same donor were harvested for further analysis (**Fig. 2A**).

**Figure 2.**
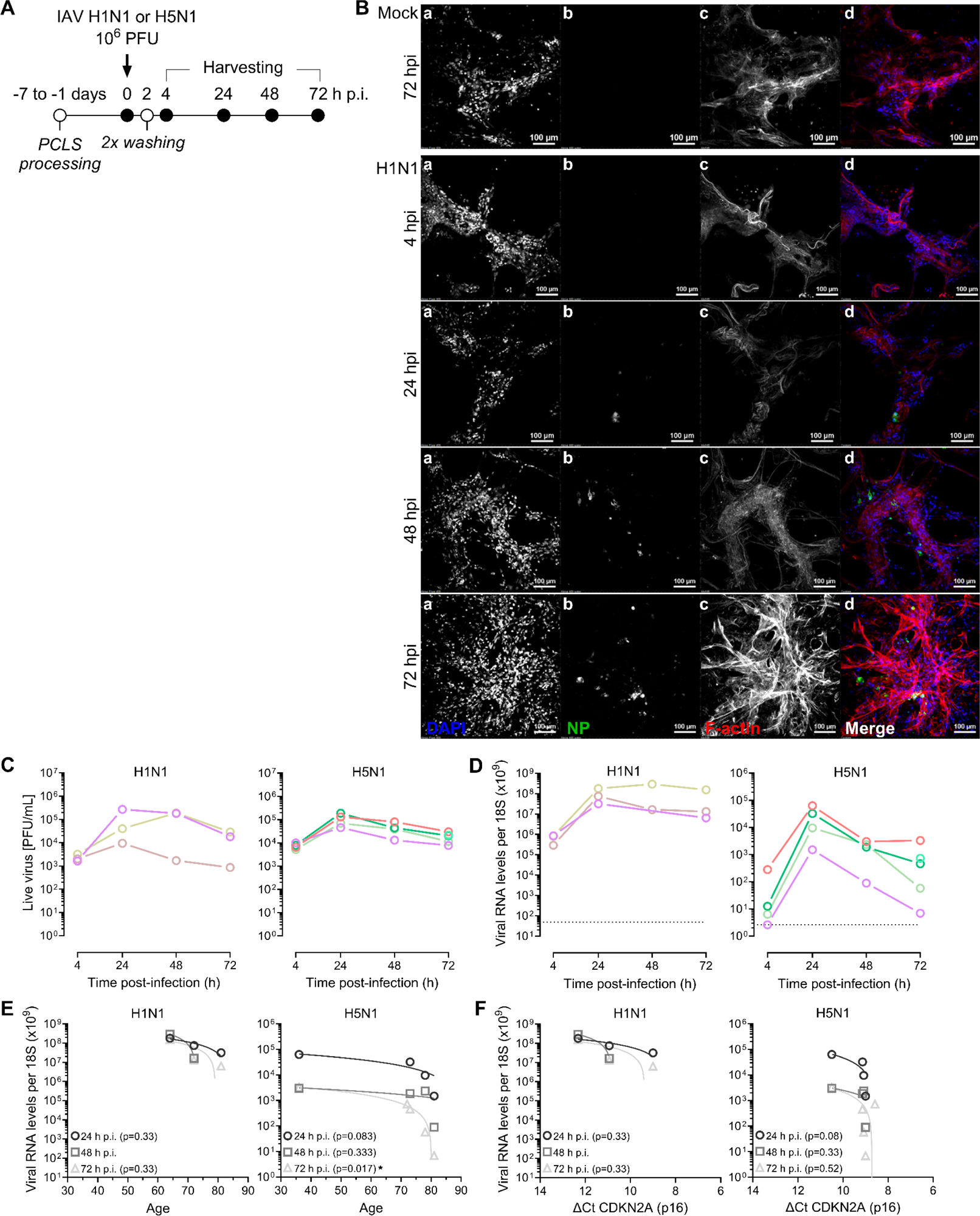
Human PCLS are productively infected by pandemic and avian IAV isolates. **(A)** PCLS were infected one to seven days after generation with 10^6^ PFU of IAV H1N1 A/HAmburg/4/2009 or H5N1 A/turkey/Turkey/2005. After 2 h PCLS were washed twice with PBS with Ca^2+^ and Mg^2+^. Growth medium was changed daily. 4, 24, 48, and 72 h p.i. supernatants and PCLS were harvested for further analysis. **(B)** Representative confocal microscopy evaluation of Mock- and H1N1-infected PCLS, 4, 24, 48, and 72 h p.i. a, DAPI; b, IAV nucleoprotein (NP); c, F-actin; d, merge: DAPI, blue; IAV NP, green; F-actin, red. Scale bars, 100 µm. **(C)** Supernatants of infected PCLS were analysed with a PFU assay. Each line represents an individual donor. **(D)** Tissue-associated viral RNA loads in PCLS over time following IAV infection expressed as IAV RNA per 18S rRNA. Dotted line represents the detection limit. **(E)** Correlation of tissue-associated viral RNA and donor age. **(F)** Correlation of tissue-associated viral RNA and deltaCT CDKN2A (p16). Association was tested using the Spearman rank correlation test. Each symbol represents an individual donor at 24 h (circle), 48 h (square), or 72 h p.i. (triangle). IAV H1N1, n=3 and H5N1, n=4.

Immunofluorescence analysis using the cytoskeleton component F-actin indicated a preserved alveolar microstructure, both in non-infected controls (Mock) and IAV-infected PCLS at all the time points p.i. analysed (**Fig. 2B**). Furthermore, the frequency of IAV NP-positive cells increased over time and individual NP-positive cells, as well as foci of infected cells, spread within the PCLS, suggesting productive IAV replication (**Fig. 2B**). Accordingly, infectious titers in the tissue culture supernatant were low at 4 hours p.i. but increased in the following time reaching a maximum of about 10^5^ PFU/ml at 24 h p.i. (**Fig. 2C**). Notably, H1N1 replicated in human PCLS in the absence of exogeneous trypsin, indicating that cellular proteases were present that cleaved the H1N1 hemagglutinin (HA) at the monobasic cleavage site (47–49). While tissue-associated viral RNA levels for H1N1 reached highest levels 24 h p.i. and remained high, viral RNA levels for H5N1 decreased about 1-2 logs from 24 h to 72 h p.i. in PCLS from all the four donors (**Fig. 2D**). Of note, RNA levels of H1N1 and H5N1 cannot be compared to each other due to distinct qPCR assays used. For further correlation analysis, we used tissue-associated viral RNA levels which, in contrast to live virus release, can be normalized to the cell number within the tissue using the housekeeping 18S rRNA. For H1N1 we could observe a trend towards lower viral RNA levels with increasing age and senescence marker CDKN2A (p16) (**Fig. 2E, F**). Furthermore, for H5N1-infected PCLS, significantly lower viral RNA levels with increasing donor age and, while not significant, as well with increasing p16 expression were measured (**Fig. 2E, F**). Altogether, our data indicate that aged/senescent PCLS are less permissive to IAV replication.

### SARS-CoV-2 infection of human PCLS is leading to a low level of replication

Next, we investigated if this age-effect towards IAV replication was also observed with another significant respiratory virus, SARS-CoV-2. Entry into target cells is mediated through binding of the viral trimeric spike protein to the cell surface receptor angiotensin-converting enzyme 2 (ACE2) after priming by transmembrane protease serine subtype 2 (TMPRSS2) (50–52). Thus, expression of ACE2 and TMPRSS2 mRNAs in PCLS was confirmed in selected donors. However, ACE2 mRNA levels were found to be lower in PCLS compared to primary human nasal cells (**Fig. 3A**). Next, using an early clinical isolate of SARS-CoV-2 (München1.1/2020/929, “Early isolate”) and the VOC SARS-CoV-2 Delta, infection experiments were performed on PCLS generated from a cohort of donors of various age (36-81 years old). PCLS were exposed to 10^6^ PFU of SARS-CoV-2 and processed as illustrated in **Fig. 3B**. Immunohistochemistry and immunofluorescence analysis revealed foci of infection at different structural localizations, suggesting a variable cell tropism of SARS-CoV-2 (**Fig. 3C, D**). In contrast to the high titers measured for IAV-infected PCLS, both SARS-CoV-2 Early and Delta isolates, replicated to lower levels, peaking at around 10^3^ PFU/ml (**Fig. 3E**). Furthermore, we assessed extracellular viral RNA of 14 PCLS donors infected with SARS-CoV-2 Early isolate and found notable levels at different timepoints p.i. (**Fig. 3F**). One donor (magenta) was negative for extracellular viral RNA during the course of infection, which was in line with undetectable infectious virus 24 h and 72 h p.i., and low titers 48 h p.i. (**Fig. 3E**). Overall, tissue-associated viral RNA levels increased during 24 h to 48 h p.i. for SARS-CoV-2 Early isolate-infected PCLS. After 48 to 72 h p.i., we could observe a plateau or slightly decreasing levels for most donors and of about one log in three out of 14 donors (**Fig. 3G**). Interestingly, within 48 h p.i. Delta isolate-infected PCLS showed an increase in tissue-associated viral RNA of about one to two logs for most donors tested. Thereafter, a plateau was reached too (**Fig. 3G**). Overall, viral RNA loads for both, the SARS-CoV-2 Early and Delta isolates, were significantly lower in aged tissue as shown in relation to donor age as well as to the expression of the senescence marker p16, indicating that aging lung cells do not provide an optimal replication environment for SARS-CoV-2 (**Fig. 3H, I**). Furthermore, association of higher ACE2 mRNA levels with increased p16 expression suggest at least partially a receptor-dependent mechanism for the observed age effect. However, ACE2 mRNA levels were independent of donor age (**Fig. 3J**). Finally, ACE2 gene expression did not change upon SARS-CoV-2 infection, an observation reported as well by others (**Fig. 3K**) (18). In summary, PCLS generated from lungs of older compared to younger individuals are less susceptible to SARS-CoV-2 infection, a feature that among other factors, might be related to the levels of the ACE2 cell-surface receptor.

**Figure 3.**
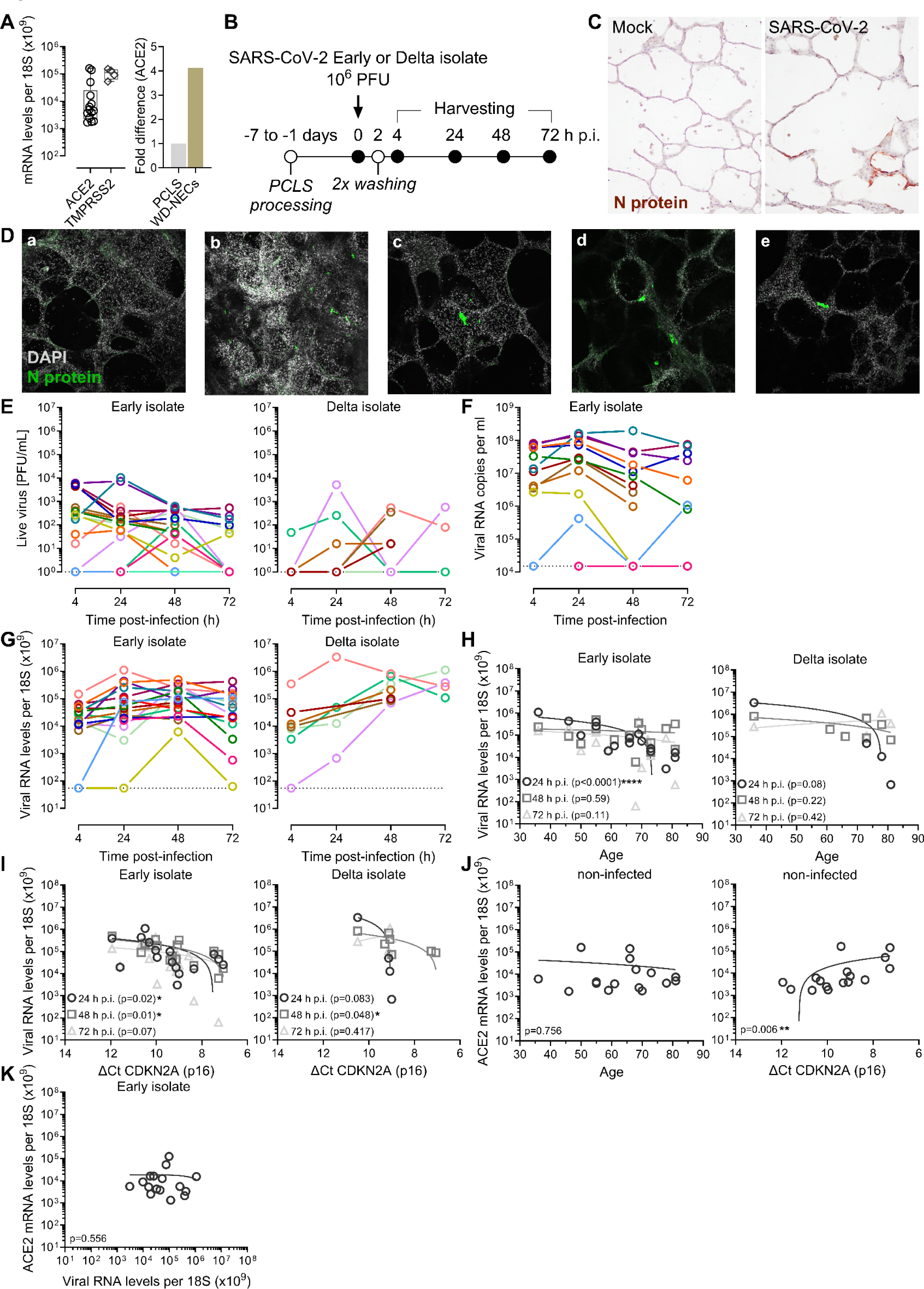
SARS-CoV-2 infection of human PCLS is leading to a low level of replication. **(A)** ACE2 and TMPRSS2 mRNA levels relative to 18S rRNA in PCLS of selected donors. Boxplots indicate the median value (centerline) and interquartile ranges (box edges), with whiskers extending to the lowest and the highest values. Each symbol represents an individual donor. (ACE2, n=15; TMPRSS2, n=4). Fold difference was calculated by comparing the median ACE2 expression of all the PCLS and WD-NEC donors tested. (PCLS, n=15; WD-NECs, n=3). **(B)** PCLS were infected one to seven days after generation with 10^6^ PFU of SARS-CoV-2 (SARS-CoV-2/München1.1/2020/929; Early isolate) and Delta isolate (Delta AY.127, hCoV-19/Switzerland/BE-IFIK-918-4879/2021). After 2 h PCLS were washed twice with PBS with Ca^2+^ and Mg^2+^. Growth medium was changed daily. 4, 24, 48, and 72 h p.i. supernatants and PCLS were harvested for further analysis. **(C)** Histological section of a representative Mock- and SARS-CoV-2 Delta isolate-infected PCLS 48 h p.i., stained for the nucelocapsid (N) protein and counterstained for haematoxylin. **(D)** Representative confocal microscopy evaluation of Mock- and SARS-CoV-2 Early isolate-infected PCLS 48 h p.i., stained for DAPI (grey) and SARS-CoV-2 N protein (green). a, Mock; b-e, SARS-CoV-2 Early isolate-infected PCLS. a-c, micrographs are projections of 3D captures. a-e, magnification 10X. **(E)** Supernatants of infected PCLS were analysed with a PFU assay. Each line represents an individual donor. SARS-CoV-2 Early isolate, n=18; Delta isolate, n=7. **(F)** Extracellular viral RNA in the supernatant of SARS-CoV-2 Early isolate-infected PCLS over time expressed as viral RNA copies per ml. Each line represents an individual donor, n=14. **(G)** Tissue-associated viral RNA loads in PCLS over time following SARS-CoV-2 infection expressed as viral RNA per 18S rRNA. Each line represents an individual donor. SARS-CoV-2 Early isolate, n=15; Delta isolate, n=7. Dotted lines represent the detection limit. **(H)** Correlation of tissue-associated viral RNA and donor age. The negative value of one donor 24 h p.i. was excluded as it was set to the detection limit of the RT-qPCR. **(I)** Correlation of tissue-associated viral RNA and deltaCT CDKN2A (p16). **(J)** Correlation of ACE2 mRNA levels in Mock-treated PCLS, 24 h p.i. and donor age or deltaCT CDKN2A (p16). Each symbol represents an individual donor. **(K)** Correlation of ACE2 mRNA and viral RNA per 18S rRNA in SARS-CoV-2 Early isolate-infected PCLS 24 h p.i. Association was tested using the Spearman rank correlation test. Each symbol represents an individual donor at 24 h (circle), 48 h (square), or 72 h p.i. (triangle).

### IAV induces cytotoxicity and a robust pro-inflammatory response in human PCLS

To investigate the impact of viral infection on the viability of lung tissue, we evaluated cytotoxicity using a LDH assay. While IAV infection led to elevated release of LDH, there was no measurable cytotoxicity upon SARS-CoV-2 infection, neither with the Early nor the Delta isolate. On the other hand, when comparing the IAV strains used, the avian H5N1 strain was clearly more cytotoxic than the H1N1 pandemic strain (**Fig. 4A, Fig. S1A**). Considering the robust inflammatory responses induced by IAV and SARS-CoV-2 infections *in vivo* (53–60), we evaluated the pro-inflammatory and antiviral responses of PCLS following infection with both viruses. To do so, we measured the mRNA levels of type I (beta) and type III (lambda 1-3) interferons (IFNs) and interleukin 6 (IL-6), as well as IFN gamma-induced protein 10 (IP10/CXCL10). Up to 72 h p.i. with SARS-CoV-2 (both Early as well as Delta isolates), there was barely any induction of IFNs detectable (**Fig. 4B, Fig. S1B, C**), most likely reflecting the low level of SARS-CoV-2 replication in PCLS (**Fig. 3E-G)**. On the other hand, H1N1-infected PCLS revealed a tendency towards higher IFN mRNA levels and avian H5N1 infection induced significant IFN responses as early as 24 h p.i. Specifically, in comparison to Mock-infected PCLS, H5N1 induced about one log, three logs, and four logs higher mRNA expression levels of IFN-β, IFN-λ1, and IFN-λ2/3, respectively (**Fig. 4B, Fig. S1D, E**). This is also in line with rapid virus replication and high viral RNA levels for H5N1 24 h p.i. (**Fig. 2C, D**). In addition, the early and remarkable antiviral response might explain the rapid decrease of H5N1 viral RNA thereafter, 48 to 72 h p.i. (**Fig. 2D**). In contrast to Mock and SARS-CoV-2 Early isolate, Delta isolate-infected PCLS tended to respond with higher IL-6 and IP10/CXCL10 mRNA levels, although not significant, most likely due to the small sample size and variability even for at Mock condition. A similar tendency in terms of IL-6 induction was observed for H1N1-infected PCLS. In contrast, H5N1 infection induced significant IL-6 expression 72 h p.i. Notably, both IAV strains led to significant induction of IP10/CXCL10 expression as early as 24 h p.i. (**Fig. 4C, Fig. S1B-E**). Altogether, SARS-CoV-2 is less cytotoxic and a weaker inducer of pro-inflammatory mediators compared to IAV H5N1 and H1N1.

**Figure 4.**
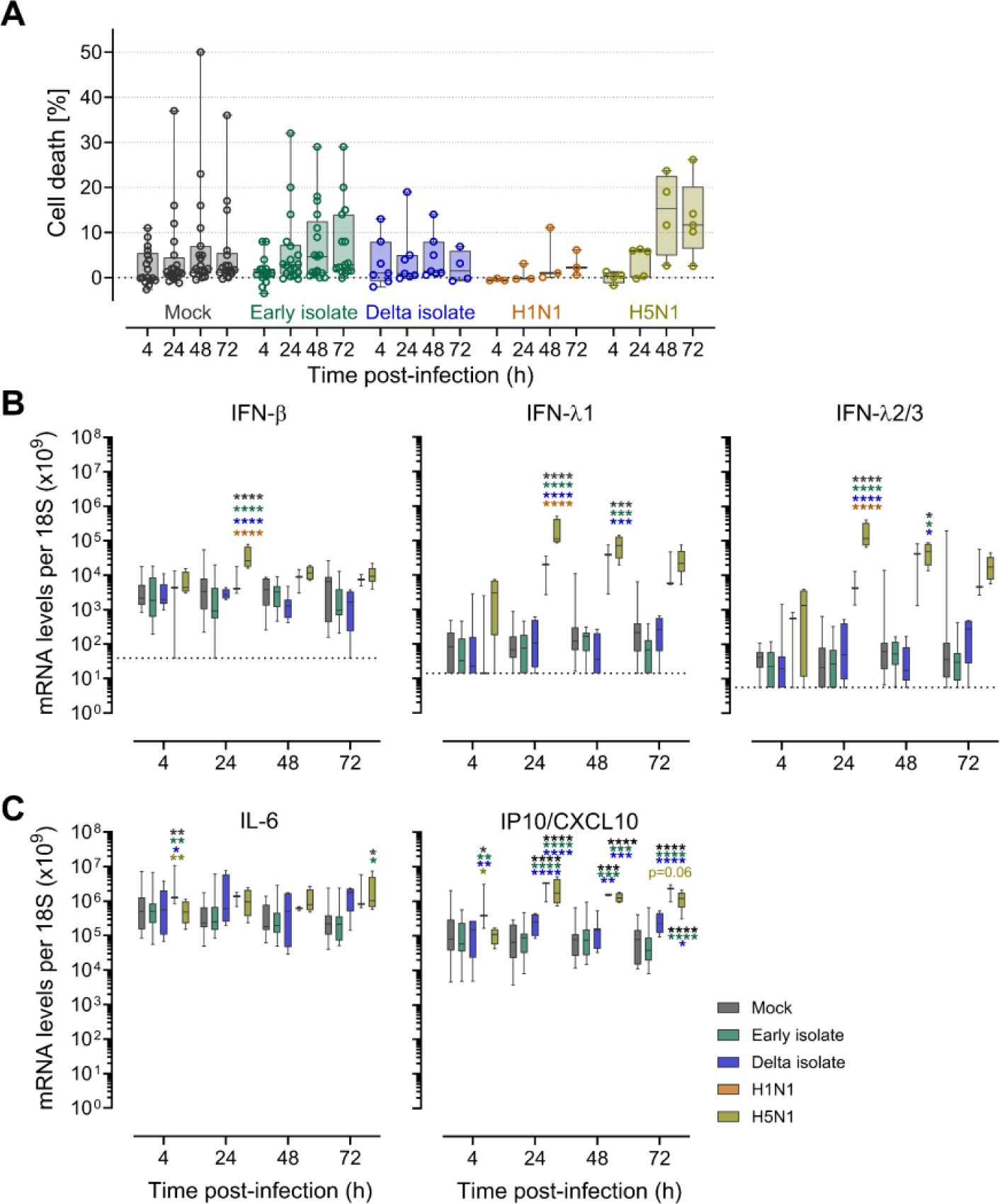
IAV is cytotoxic and induces a robust pro-inflammatory response in human PCLS. **(A)** Cell death (%), calculated from lactate dehydrogenase (LDH) release in PCLS, exposed to 10^6^ PFU of SARS-CoV-2 Early or Delta isolate, IAV H1N1, or H5N1 over time. Each symbol represents an individual donor. **(B)** IFN-β, IFN-λ1, IFN-λ2/3 and **(C)** IL-6, IP10/CXCL10 mRNA expression levels per 18S rRNA over time. A two-way ANOVA and the Tukey’s multiple comparisons test was applied to compare the different groups. Mock, black; SARS-CoV-2 Early isolate, green; SARS-CoV-2 Delta isolate, blue; IAV H1N1, orange; IAV H5N1, yellow. Boxplots indicate the median value (centerline) and interquartile ranges (box edges), with whiskers extending to the lowest and the highest values.

## DISCUSSION

In this study, we explored the impact of donor age on the susceptibility of the human distal lung epithelium to respiratory virus infection. To do so, we used an *ex vivo* model based on PCLS cultures prepared from lung tissue of individuals spanning from 36 to 81 years of age. Our data demonstrate that donor age is associated with distinct susceptibility of PCLS to IAV and SARS-CoV-2 infection, with PCLS from older versus younger patients being less permissive to both viruses. Furthermore, while SARS-CoV-2 infection was leading to undetectable cell death and triggered only limited host responses, IAV infection caused significant cytotoxicity and pro-inflammatory responses.

The high and low productive replication potentials in the distal lung for several IAV strains and SARS-CoV-2 variants, respectively, have been described before using lung explant models (16–18). When comparing donors of different age, PCLS in contrast to lung explant cultures ensure better homogeneity and reproducibility and reduce the degree of variability between sample replicates, due to precise cutting and because consecutive slices can be obtained from the same lung explant. While SARS-CoV-2 Early and Delta isolates replicated at low levels in PCLS, IAV H1N1 and H5N1 replication resulted in the release of significant numbers of infectious viral particles into the culture supernatant. The differential susceptibility of PCLS to SARS-CoV-2 and IAV infection is possibly due to differences in the alveolar cell surface expression of cognate virus receptors. For instance, ACE2 expression was lower in PCLS compared to WD-NEC cultures and has been shown to gradually decrease from the upper to the lower human respiratory tract and thereby relating to virus infectivity (61). A similar coherence of receptor expression levels and susceptibility to SARS-CoV-2 infection was recently reported for human placenta explants (62). Furthermore, using a high-sensitivity RNA *in situ* mapping approach, ACE2 was revealed to be expressed in less than 1% of alveolar type 2 cells (61, 63, 64). In alveolar organoids, Hönzke *et al.* showed limited permissiveness of the human distal lung for SARS-CoV-2 infection due to sparse ACE2 levels and that viral replication can be increased by ACE2 overexpression (18). Nevertheless, SARS-CoV-2 infection of the lower respiratory tract was shown in pathological samples of COVID-19 patients (65, 66). The virus likely reaches the distal lung via droplets aspiration from the naso- and oro-pharynx, although the exact mechanisms are unknown. In contrast, the HA of IAV primarily binds to sialic acid residues (SA, *N*-acetylneuraminic acid), that are part of cell surface glycoconjugates which mediate virus attachment to the host cell (67). The binding of human and avian IAV to SA in α2,6- and α2,3-glycosidic linkage, respectively, is believed to be an important species barrier (68–70). While SAα2,6 residues are abundantly expressed on airway cells throughout the conducting airways (71, 72), SAα2,3 residues are predominantly expressed by alveolar type 2 cells and resident macrophages of the distal lung and less abundantly expressed in the human upper respiratory tract. Accordingly, avian IAV were shown to replicate efficiently in the lower respiratory tract while replication in the upper respiratory tract is very limited (68–70). Consequently, human-to-human transmission of avian IAV such as the highly pathogenic H5N1 is rare (73). Still, H5N1 IAV is a prevailing pandemic threat and led to 882 confirmed cases between 2003 and 2023, with a case fatality rate as high as 52% (74).

Interestingly, for both viruses tested, we found an association of lower viral RNA levels with older age of the tissue donor and with higher CDK2NA (p16) expression levels. Generally, association of viral load dynamics with age is highly debated in literature. Whereas in some studies no differences in SARS-CoV-2 RNA loads between children and adults were found (75, 76), others reported slightly lower viral RNA loads in children (77, 78). ACE2 expression was shown to be age-dependent in the nasal epithelium (79) as well as in alveolar type 2 cells (80), with lower expression in children compared to adults. However, our study focused on lung tissue from adult donors and didn’t include younger adults and children. In another study no differences in ACE2 transcript levels in lung samples from deceased humans of different age were found (81). Furthermore, senescence has been reported to lead to enhanced expression of ACE2 in primary human small airway epithelial cells, when treated with condition media of senescent cells (13). In line, we found that higher ACE2 expression correlated with higher CDK2NA (p16) expression levels. Since senescent cells accumulate with age and have other potentially adverse functions, this phenomenon might explain why the elderly are more susceptible to severe COVID-19 (82). However, the interplay between viruses and cellular senescence is complex. In some cases, virus infection can induce cellular senescence and subsequently restrict or improve virus propagation, while in others, certain viruses evolved strategies to subvert senescence (83). In contrast to a previous study on IAV-induced senescence (12), we observed lower viral loads in the aged pulmonary tissue for all the IAV strains and SARS-CoV-2 isolates tested. Nevertheless, using comprehensive approaches to analyse autopsy-derived lung specimens of patients with COVID-19 in contrast to age-matched control individuals, Wang *et al.* identified parenchymal lung senescence as a contributor to COVID-19 pathology (84). Thus, illustrating the limitations of *ex vivo* models, acute infection of PCLS might not reflect the effect of virus infection on senescence and vice versa. However, specific characteristics of senescence are cell type specific and thus would require in-depth analysis using for example single-cell RNA sequencing, proteomics, or related approaches (85, 86), whereas many of those are already established for the PCLS model (87).

Several studies have demonstrated that H5N1 infection leads to a more robust induction of cytokines and chemokines compared to H1N1 (88–92). Furthermore, H5N1 infected humans present unusually high serum concentrations of pro-inflammatory cytokines and chemokines, which are believed to contribute to disease severity (57–59, 93). Similarly, characteristic features of severe COVID-19 rely in the excessive release of pro-inflammatory cytokines, a so-called “cytokine storm”, and an immune dysregulation, specifically, a massive immune stimulation resulting in a macrophage activation syndrome, aberrant neutrophil activation, or T cell hyperreactivity. Notably, the most destructive phase of immune-driven pathogenesis often happened when the viral RNA was no longer detectable (53, 94–97). However, our experimental set up only reflects the acute phase of infection and not what happens *in vivo* one to two weeks post infection. SARS-CoV-2 infection of PCLS did not lead to significant induction of IL-6 or IP10/CXCL10 expression, indicating that the local cellular environment alone, independent of circulating immune cells, cannot cause the massive local cytokine production seen in severe COVID-19 cases. On the other hand, H5N1 and to lesser extent H1N1 induced significant IL-6 and IP10/CXCL10 expression and high cytotoxicity levels, suggesting a contribution of the alveolar epithelium in the overexuberant inflammatory responses observed in severe IAV cases. The differences in host response observed between SARS-CoV-2- and IAV-infected PCLS likely also reflect the distinct viral replication profiles.

Additionally, our results suggest that the pulmonary environment remains immunocompetent throughout all ages tested. Even though there are some physiological limitations, the PCLS model is obviously purposeful to analyse certain questions independent of circulating immune cells.

In summary, our data show that pulmonary tissues from older donors are generally less susceptible to IAV and SARS-CoV-2 replication compared to tissues from younger donors. In contrast to SARS-CoV-2, IAV replicated more efficiently in the human alveolar epithelium leading to more pronounced innate immune responses. Our findings indicate that the elderly are not more susceptible to respiratory virus infection than young people as a rational of elevated risk for severe disease by local viral replication only, but points towards other mechanism such as immune-mediated pathogenesis.

## ACKNOWLEDGMENTS

This work acknowledges support from the Lungenliga Bern, Switzerland to M.P.A. (Project ID “The impact of SARS-CoV-2 infection on the aged lung”), and by intramural funding of the Institute of Virology and Immunology, Switzerland. The funders had no role in study design, data collection and analysis, decision to publish, or preparation of the manuscript. The authors would like to thank all study participants and their families.

## AUTHOR CONTRIBUTIONS

MFC and MPA conceived and designed research; MB, CM, BZ, CC, DJ, PD, TMM performed experiments; MB, CM, BZ, CC, MFC, MPA analyzed data and interpreted results of experiments; MB, CM, BZ, and CC prepared figures; MB, CM, MFC, MPA drafted manuscript; MB, CM, BZ, CC, GZ, VT, MFC, MPA edited and revised manuscript; all authors approved final version of manuscript.

---

**Fig. S1:**
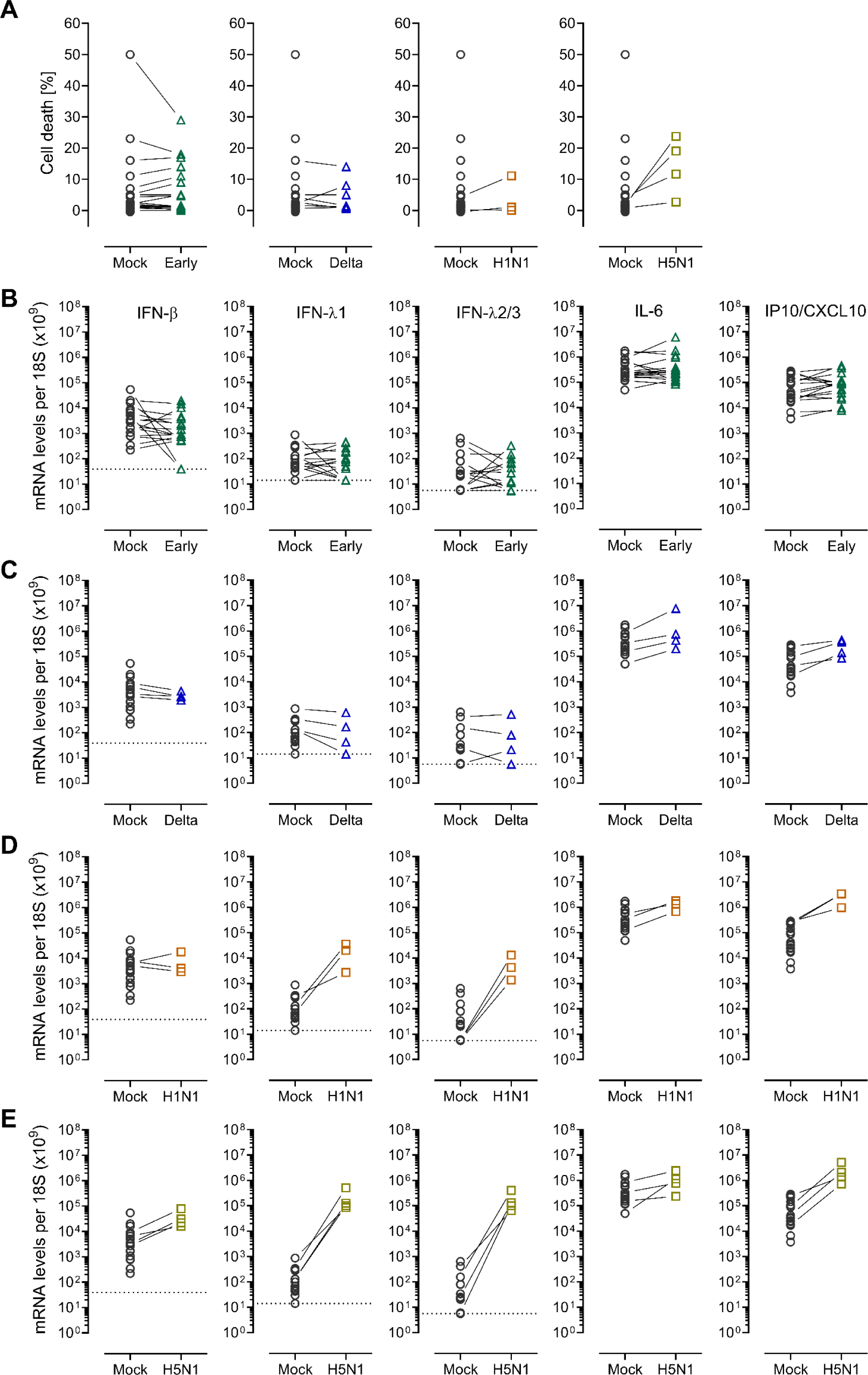
Representation of cell death and host response per donor. **(A)** Cell death (%), calculated from lactate dehydrogenase (LDH) release in PCLS, exposed to 10^6^ PFU of SARS-CoV-2 Early or Delta isolate, IAV H1N1, or H5N1 48 h p.i. Each symbol represents an individual donor, whereas lines connect Mock and virus conditions of the same donor. **(B, C, D, and E)** IFN-β, IFN-λ1, IFN-λ2/3, IL-6, and IP10/CXCL10 mRNA expression levels per 18S rRNA 24 h p.i. Each symbol represents an individual donor, whereas lines connect Mock and virus conditions of the same donor for **(B)** SARS-CoV-2 Early isolate-, **(C)** Delta isolate-, **(D)** IAV H1N1-, and **(E)** H5N1-infected PCLS. Mock, black; SARS-CoV-2 early isolate, green; SARS-CoV-2 Delta isolate, blue; IAV H1N1, orange; IAV H5N1, yellow.

